# Modeling Dyslexia in Neurotypical Adults by Combining Neuroimaging and Neuromodulation Techniques: A Hypothesis Paper

**DOI:** 10.1101/2025.05.04.651804

**Authors:** Daniel Gallagher, Zian Huang, Shinri Ohta

## Abstract

Dyslexia is a prevalent developmental disorder marked by deficits in literacy skills. Given that the core deficits of dyslexia are uniquely human, animal models have not been as useful in dyslexia research as they have been in other areas of research. While significant progress has been made through behavioral and neuroimaging studies, a viable model could facilitate controlled investigations into the neural mechanisms underlying dyslexia and accelerate the development of targeted interventions. In this hypothesis article, we propose a two-pronged approach to model dyslexia in neurotypical adults using neuroimaging and neuromodulation techniques. First, we propose using functional and structural MRI data to cluster individuals into neuropathologically derived subgroups in order to facilitate the classification of dyslexia subtypes based on neuropathological characteristics. Second, we propose employing transcranial temporal interference stimulation (tTIS) to temporarily downregulate activity in brain regions specified in the clustering analysis, inducing subtype-specific dyslexic symptoms in neurotypical individuals. This approach enables the establishment of causal or probabilistic relationships between neuropathologies and dyslexia subtypes, while at the same time creating dyslexia models to facilitate investigation into subtype-specific interventions. By integrating neuroimaging and neuromodulation, we hope to offer a viable substitute for animal models in dyslexia and accelerate the development of personalized therapeutic strategies for dyslexia.

## 1 Introduction

As a learning disability causing literacy skill deficits, dyslexia, which includes several subtypes, is one of the most prevalent developmental disorders affecting the human population, affecting about 7% of the population (Yang et al., 2022). Dyslexia is highly heterogeneous, with different individuals exhibiting distinct cognitive deficits. Broadly, subtypes have been identified based on impairments in phonological awareness (PA), rapid automatized naming (RAN), and visuo-spatial processing, among other cognitive domains (Bruck, 1992; Denckla & Rudel, 1976). As the heterogeneity of dyslexia has come into greater focus, models have gradually shifted from emphasizing single deficits to double deficits to multiple deficits (Pennington, 2006; Wolf & Bowers, 1999). Although the multiple deficit model provides a strong explanatory framework for understanding its heterogeneity (McGrath, Peterson, & Pennington, 2020; Pennington, 2006), the most appropriate approach to classifying dyslexia subtypes remains an open debate.

Some neuroimaging research has provided significant insights into the neurobiological basis of dyslexia subtypes, highlighting structural and functional differences in key reading-related brain regions. For example, hypoactivation, atypical connectivity patterns, and structural variations in gray matter volume have all been reported to vary according subtype (Jednoróg, Gawron, Marchewka, Heim, & Grabowska, 2014; Norton, Beach, & Gabrieli, 2015). Despite these advances, most studies investigating the neural mechanisms of dyslexia do not account for specific behavioral deficits, representing a gap in the dyslexia literature.

Beyond the outstanding need to more fully clarify the neurobiological basis of dyslexia subtypes, the development and testing of new dyslexia treatments remains hindered by the need to recruit sufficiently many participants with the specific subset of deficits of interest. Other neurodevelopmental and neurological disorders, such as attention-deficit hyperactivity disorder (ADHD), autism, etc., mitigate this challenge by utilizing appropriate animal models (Russell, 2011; Varghese et al., 2017), which replicate in an animal the symptomatic expressions of the disorder being studied. For example, rats may be selectively bred to achieve specific symptoms, as is done with spontaneously hypertensive rats in ADHD research (Sagvolden et al., 1992). In other cases, the gene of a mouse are modified (e.g., knockout and knockin mice), and still in other cases, drugs may be administered to induce the desired symptoms. Although not without their limitations (Robinson et al., 2019), animal models serve as an important investigative tool that allow researchers to delineate neuropathologies and assess the viability of various therapeutic and pharmacological interventions (Mukherjee, Roy, Ghosh, & Nandi, 2022). These models allow researchers to do this substantially faster than would otherwise be possible solely using human subjects, ultimately shortening the time it takes for novel interventions to reach clinical implementation.

However, unlike other neurological disorders, dyslexia specifically affects literacy skills, which are uniquely human. Therefore, an animal model of dyslexia cannot adequately replicate the challenges faced by human individuals with dyslexia. Even if, for example through genetic modification, “dyslexia” were to be artificially induced in an animal, it is unclear how it should manifest and what symptoms should present. Nevertheless, the approach has been taken in genetic studies of dyslexia. In one study, KIAA0319 knockdown rats—where the KIAA0319 gene was suppressed by RNA interference—presented with impaired phoneme processing in the primary auditory cortex (Centanni et al., 2014). In another genetic study, DCDC2 knockdown rats presented with speech sound discrimination deficiencies (Centanni et al., 2016). These and similar studies offer crucial insights into the multifactorial genetic underpinnings of dyslexia and are indeed important pieces of the puzzle. However, because animals are incapable of literacy, it is not possible to surmise whether the induced symptoms truly reflect dyslexia or just peripherally related symptoms, and by extension therefore, it is not possible to assess the extent of applicability of the models to human dyslexia.

Dyslexia researchers are therefore faced with a unique challenge, where they must either rely on genetically modified animals with an unknowable degree of symptomatic specificity or work exclusively with dyslexic individuals, which necessarily entails higher hurdles related to recruiting, turnaround times, interindividual differences, etc. (Roitsch & Watson, 2019). Thus, a reliable model of dyslexia would fill a gap in investigatory approaches in dyslexia research. However, again, since literacy skills—the principal marker of dyslexia—are unique to humans, an animal model is simply not feasible. We therefore propose the development of a dyslexia model in neurotypical adult humans.

To develop a dyslexia model in neurotypical adults, we put forward a two-part hypothesis. Namely, we hypothesize that 1) developmental dyslexia leads to anatomical and functional abnormalities at specific regions in the adult brain differentiated by the subtype of dyslexia, and 2) by using transcranial temporal interference stimulation (tTIS) on those target regions, we can temporarily induce the functional deficiencies that give rise to subtype-specific symptoms of dyslexia. Correspondingly, we take a two-pronged approach to developing the model. In the first prong, we establish the neurological basis of dyslexia by analyzing open-source brain data on adults and children with and without dyslexia in order to elucidate the most relevant brain abnormalities associated with the disorder and select target regions. In the second prong, we employ tTIS, which can achieve both focal and deep brain stimulation without stimulating surrounding areas, at the target regions specified by the structural and functional analysis to induce subtype-specific dyslexic symptoms in neurotypical adults. In other words, we expect that by stimulating a given brain region associated with a given subtype of dyslexia, only the symptoms of that subtype should be elicited. In this way, subtype-specific dyslexia models can be created in neurotypical adults, facilitating the development of more targeted and individualistic interventions for treating dyslexia.

## 2 Background

### 2.1 Dyslexia Subtypes

Dyslexia in the broad sense is behaviorally characterized by a deficiency in literacy skills, such as lower reading accuracy or fluency, without affecting general intelligence or other linguistic abilities. The most prominent region employed in the reading network is well-known to be the left fusiform gyrus (FG), also known as the visual word form gyrus (VWFA) (Cohen & Dehaene, 2004; Cohen et al., 2000). For example, it has been shown that reading speeds positively correlate with the degree of VWFA activation (Christodoulou et al., 2014; Langer, Benjamin, Minas, & Gaab, 2015). More broadly, however, the cognitive act of reading consists of both cooperative and competitive mechanisms recruiting many areas involved in orthographic, phonological, and semantic processing, such as the left inferior frontal gyrus (LIFG) and left temporal, left inferior parietal, and occipito-temporal regions (Cattinelli, Borghese, Gallucci, & Paulesu, 2013).

Various regions in the reading network have been associated with both structural and functional abnormalities in dyslexia. For example, a meta-analysis of voxel-based morphometry (VBM) studies showed that individuals with dyslexia exhibited gray matter reduction in the right superior temporal gyrus, left superior temporal sulcus, while the VWFA was shown by several individual studies to have gray matter reduction without meeting clustering criteria for the meta-analysis. (Richlan, Kronbichler, & Wimmer, 2013). Another meta-analysis focusing on functional abnormalities revealed consistent hypoactivation in the left inferior parietal lobule, LIFG, superior temporal, middle temporal inferior temporal and fusiform regions (e.g., VWFA), as well as hyperactivation of the primary motor cortex and anterior insula (Richlan, Kronbichler, & Wimmer, 2009). However, as a neurodevelopmental disorder, it makes intuitive sense that dyslexia should affect individuals differently at different stages of development. Indeed, when controlling for age group, the results diverged slightly. It was found that while both children and adults with dyslexia exhibited hypoactivation in the left ventral occipital-temporal region (which includes the VWFA), only children exhibited hypoactivation in bilateral inferior parietal regions, while only adults showed hypoactivation in the superior temporal regions (Richlan, Kronbichler, & Wimmer, 2011).

These findings have already clarified a great deal of the neurobiology of dyslexia. Nevertheless, most neuroimaging studies of dyslexia are confounded by the heterogeneity of the disorder, which can be partially alleviated by appropriately classifying individuals according to their specific deficit(s). Thus, it is necessary to identify the distinctive neural bases of the different dyslexia subtypes. To that end, Norton and colleagues (2015) neatly summarized the contemporary understanding of the brain bases of behaviorally derived phenological subtypes of dyslexia. Based on their findings, some core behavioral deficits associated with dyslexia are recognized: Phonological awareness (PA), rapid automatized naming (RAN), and sensory and working memory related processes (Norton et al., 2015).

In the phonological deficit hypothesis of dyslexia, it is thought that dyslexia is the result of poor phonological skills hindering the acquisition of the rules governing spelling (M. Snowling, 1998). These PA deficits manifest in behavioral experiments as impaired repetition and decoding of nonwords (Rack, Snowling, & Olson, 1992; M. J. Snowling, 1981) and impaired recognition of rhymes and alliterations (Bradley & Bryant, 1978). Such deficits have been shown to arise from functional and structural connectivity (as measured by diffusion tensor imaging) between auditory cortices and the LIFG, reduced prefrontal activation, but no abnormalities in temporal lobe activation (Boets et al., 2013).

In RAN deficits, individuals with dyslexia exhibit markedly slower naming speeds for colors, numbers, letters, and objects (Denckla & Rudel, 1976). These deficits are less localized than PA deficits and are associated with more whole-brain volumetric differences (He et al., 2013), as well as lower activation in the right cerebellar lobule VI (Norton et al., 2014).

Some studies have also shown that individuals with dyslexia may have various abnormalities in sensory and working memory related processes, such as reduced left-lateralized entrainment at frequencies critical for parsing speech signals (Giraud & Poeppel, 2012; Lehongre, Morillon, Giraud, & Ramus, 2013), reduced left-lateralized integration of phonological and orthographic information (Hasko, Bruder, Bartling, & Schulte-Körne, 2012), and reduced bilateral activation in the BA7 leading to a working memory deficit related to temporal order processing (Beneventi, Tønnessen, & Ersland, 2009).

In one study specifically comparing various dyslexia subtypes, Jednoróg and colleagues (2014) categorized dyslexic children into subtypes based on behavioral assessments, including PA, RAN, and sensory deficits, and found specific gray matter patterns aligning with the dyslexia subtypes. Specifically, their voxel-based morphometry (VBM) approach revealed the LIFG, cerebellum, right putamen, and bilateral parietal cortex as areas with gray matter volume differences between the different dyslexic subtypes (Jednoróg et al., 2014).

### 2.2 The Multiple Deficit Model of Developmental Disorders

Thus far, we have discussed dyslexia as though there is a one-to-one correspondence between a given brain anomaly and a specific behavioral symptom. However, the multiple deficit model (MDM) has gained prominence in recent years and challenges this notion by proposing that developmental disorders, including dyslexia, arise from the interactive effects of multiple risk factors rather than a single causal deficit (Pennington, 2006). Unlike single-deficit and double-deficit models, which assume that a particular brain anomaly leads directly to a specific cognitive impairment, the MDM conceptualizes dyslexia as a probabilistic outcome resulting from the accumulation and interaction of multiple neural, genetic, and environmental influences.

From this perspective, a given brain anomaly does not necessarily and deterministically produce a specific deficit but rather increases the probability of it. Conversely, the same cognitive symptom can arise from different underlying neural anomalies in different individuals. This model has demonstrated more reliable predictive power than single-deficit models, particularly for individuals with dyslexia and dyscalculia, though a hybrid approach using the different models in tandem seems to outperform using either model exclusively (McGrath et al., 2020; Pennington et al., 2012). Importantly, within this framework, the discussion of deficits and subtypes turns from “causally deterministic” to “probabilistically predictive,” and rather than speaking of “core deficits” of dyslexia (such as PA and RAN deficits), the more appropriate terminology is “predictors.”

Nevertheless, even assuming MDM and discarding the notion of deterministic relationships between specific brain regions and symptoms, studying the likelihood of specific neural anomalies contributing to particular cognitive deficits remains crucial for understanding the neurobiology of dyslexia and guiding the development of effective interventions. By identifying which brain regions are most likely to contribute to specific deficits, we can refine dyslexia diagnosis and develop more targeted therapies that address the individualized constellation of risk factors present in each case.

### 2.3 Defining the Neurological Pathogenesis of Dyslexia Subtypes

Of the neuroscientific studies targeting dyslexia subtypes, most have started with the symptomatic manifestations of dyslexia and then aimed to elucidate the neural patterns associated with those symptoms. However, due to the complex etiology of dyslexia, it is challenging to fully describe the neural basis of specific dyslexia subtypes using only monomodal analyses. To solve this challenge, we propose grouping individuals with dyslexia based on similarities in their structural and functional MRI data in order to uncover novel, neuropathologically defined subtypes. This represents a new perspective, contrasting with previous research approaches that typically focus on grouping dyslexia by specific behavioral patterns or symptomatic manifestations. By emphasizing the identification of subtypes through structural and functional brain data, we expect to provide new insights for targeted interventions tailored to each subtype of dyslexia.

By combining this approach with tTIS, we suggest that the relationships between the implicated brain regions and specific dyslexia symptoms can be established, and dyslexia models can be created in neurotypical adults. Previous studies employing other forms of non-invasive brain stimulation (NIBS), such as transcranial direct current stimulation (tDCS) and transcranial magnetic stimulation (TMS) have already demonstrated stimulation at various sites as an effective therapeutical intervention regardless of age group, though the pressing need for more NIBS studies is acknowledged (Turker & Hartwigsen, 2022). Towards that goal, tTIS in particular can fill an important gap in the dyslexia-NIBS literature thanks to its unique capability to stimulate previously inaccessible regions, including the VWFA.

The aforementioned dysfunction observed in the VWFA, LIFG, cerebellum (lobule VI), and superior temporal regions suggest the viability of targeting these regions for localized brain stimulation in adults. However, by additionally looking at clusters of neuropathologies found in individuals with dyslexia, other regions of interest may be observed and subsequently tested via tTIS. In the subsequent sections, we will describe the specific methodologies and benefits of our two-pronged approach to developing a dyslexia model in neurotypical adults.

## 3 Functional and Structural MRI Analysis

The first prong of our approach aims to identify the brain regions associated with dyslexia subtypes by employing two complementary methods to assess brain anomalies: structural magnetic resonance imaging (MRI) to examine brain anatomy and functional MRI (fMRI).

By analyzing both structural and functional data, it is possible to determine how structural changes correlate with functional connectivity patterns. This could lead to a more nuanced understanding of dyslexia and provide valuable information for intervention strategies. Future avenues can explore more detailed functional connectivity analysis or the use of other brain atlases and ROIs to refine our understanding of the relationship between brain structure and function in dyslexia.

### 3.1 Structural MRI Analysis

By performing voxel-based morphometry (VBM), and surface-based morphology (SBM), it is possible to elucidate significant structural differences between individuals with dyslexia and controls. VBM analysis focuses on the 3D volume of brain tissue and is primarily used for analyzing gray and white matter volumes across brain regions; while SBM works with the cortical surface, analyzing features such as cortical thickness, sulcal depth, and gyrification, and is often used to study more localized cortical regions (Goto et al., 2022).

Through VBM, it is possible to assess whether dyslexic adults exhibit reductions in grey/white matter volume in regions crucial for language processing and visual-spatial integration. For instance, reductions in the volume of key white matter tracts, such as the arcuate fasciculus (Žarić et al., 2018) and the inferior fronto-occipital fasciculus (Lou et al., 2019), may indicate decreased efficiency in neural connectivity, potentially reflecting challenges in recruiting neural resources. Additionally, reductions in the left corpus callosum may disrupt interhemispheric communication, impacting functions commonly associated with the right hemisphere (Hynd et al., 1995).

Moreover, SBM analysis uniquely allows for the measurement of cortical thickness, enabling the detection of subtle structural changes that may not be captured by other voxel-based methods. Through SBM, cortical thinning can be identified in key regions such as the left fusiform gyrus (FG), superior temporal gyrus (STG), middle temporal gyrus (MTG), and inferior frontal gyrus (IFG) (Williams, Juranek, Cirino, & Fletcher, 2018).

While VBM and SBM work as whole-brain analysis requiring correcting for thousands of voxels/vertices (e.g., FWE or FDR correction), making it harder to find significant effects, ROI-based analysis limits comparisons to a smaller number of voxels/vertices, which will lower the correction burden. However, it also relies on prior knowledge to define ROIs, which may introduce bias and limit the discovery of unexpected findings. Combining whole-brain and ROI-based approaches can thus provide a more comprehensive understanding of dyslexia-related structural alterations.

Notably, cortical thinning in these regions might be more pronounced in older adults with dyslexia, potentially reflecting an age-related pattern that offers valuable insights into the evolution of dyslexia across the lifespan. Furthermore, it is promising to investigate whether there might be differential aging patterns between the left and right hemispheres, as compensatory mechanisms could be at play in dyslexic brains. These findings could help guide the selection of optimal stimulation sites for targeted interventions aimed at enhancing dyslexia-related cognitive functions.

However, since brain stimulation can modulate neural activity, relying solely on structural results does not offer a robust or reliable foundation for defining stimulation targets. Therefore, in our future analyses, integrating both structural and functional MRI findings to identify areas of overlap between the two will yield more meaningful insights.

For a preliminary structural analysis based on the described methods, see the supplementary file, which includes data from the preliminary analyses conducted with the datasets from Banfi et al. (2021) and Cavalli, Chanoine, & Ziegler (2023).

### 3.2 Functional MRI Analysis

To investigate the neural mechanisms underlying dyslexia, we hypothesize that functional differences between individuals with dyslexia and typical readers are associated with structural brain alterations in dyslexic individuals. To test this hypothesis, we will employ a comprehensive analysis of both functional and structural MRI data.

Given that the research of both data sets focuses on the difference in functional activity between the dyslexic group and typical readers, we expect to first replicate findings from previous studies to confirm the existence of functional differences. Additionally, by integrating the observed structural difference in dyslexic brains with the distinct functional patterns associated with dyslexia, it is possible to further explore if their functional patterns contribute to or reflect structural changes.

Secondly, to effectively investigate the efficacy of dyslexia interventions, it is crucial to gain a comprehensive understanding of the brain networks and pathologies underlying the condition. Using generalized Psychophysiological interactions (gPPI), it is possible to characterize task-related modulation to reveal specific changes in whole-brain connectivity between subject groups (McLaren, Ries, Xu, & Johnson, 2012). Additionally, by computing ROI-to-ROI connectivity (RRC) as outlined by Nieto-Castanon & Whitefield-Gabrieli (2022), we can examine functional connectivity between regions of interest (ROIs) and investigate how these patterns differ between dyslexic and typical readers. Previous research has shown that disrupted network interactions serve as a neural marker for dyslexia, with dyslexic individuals exhibiting abnormal task-related functional connectivity that negatively impacts reading performance (Turker, Kuhnke, Jiang, & Hartwigsen, 2023).

Through this methodology, we aim to enhance our understanding of the neural mechanisms underlying dyslexia, specifically how functional connectivity may relate to structural alterations in key brain regions. If any regions show alignment between functional and structural results, they will be promising targets for transcranial temporal interference stimulation (tTIS). Regions exhibiting both structural alterations (e.g., reduced cortical thickness or volume) and functional connectivity disruptions may indicate core neural deficits underlying dyslexia. These areas are likely to play a crucial role in reading-related processing, making them potential candidates for targeted neuromodulation.

### 3.3 Principal Component Analysis

The variability observed in our preliminary results of structural data (see supplementary materials) still showed a larger variance than controls, reflecting the heterogenous nature of dyslexia and suggesting the possibility that the data sets combined different subtypes of dyslexia. Therefore, we plan to apply principal component analysis (PCA) to reduce the dimensionality of structural and functional MRI data and identify the principal components that best explain differences between dyslexic subtypes. PCA simplifies complex data by extracting the majority of variance (see, e.g., Stoyanov et al., 2019).

We first propose that PCA can be used to identify brain regions with the most significant deviations in characteristics such as cortical thickness, volume, and density. These deviations may correspond to distinct brain structures associated with dyslexia subtypes, independent of symptoms. Rather than categorizing participants by pre-defined subtypes, PCA will group participants based on structural anomalies revealed by the data itself, offering a data-driven approach.

To refine the analysis, Varimax rotation can enhance the clarity of the extracted components. This method ensures that each brain region strongly loads onto only one component, revealing clearer structural patterns (Van Boxtel, 1998). After rotation, factor loadings will show how each component contributes to structural variability, while factor scores will reflect how these components relate to individual participants. These scores can be analyzed to see if structural differences vary systematically between dyslexic and control groups, providing insight into potential neuroanatomical subtypes of dyslexia.

Beyond structural analysis, PCA can be extended to functional connectivity data to explore whether the identified subgroups also exhibit distinct functional connectivity patterns. This step will help determine if structural differences correspond to disruptions in large-scale brain networks involved in reading and language processing. PCA will be applied to the functional connectivity matrix to identify principal components that explain the variance in connectivity patterns caused by potential dyslexia subtypes. These subtype-specific patterns form the foundation of neuroscientific diagnostics by linking structural and functional data to functional impairments, ultimately contributing to more personalized and targeted interventions.

## 4 Transcranial Temporal Interference Stimulation (tTIS) for Modeling Dyslexia Subtypes

The second prong of our approach involves using tTIS to stimulate the various regions highlighted by the functional and structural analysis. One major category of NIBS is transcranial electrical stimulation (tES).

### 4.1 Non-invasive Brain Stimulation (NIBS)

Multiple NIBS methods have shown great utility in cognitive neuroscientific studies, especially for establishing causal relationships between brain regions and cognitive functions (Bergmann & Hartwigsen, 2021) and for providing novel and promising therapeutic intervention for disorders like autism and dyslexia (Lazzaro et al., 2022; Sokhadze et al., 2014; Turker & Hartwigsen, 2022). For example, transcranial direct current stimulation (tDCS) has revealed the causal role of the LIFG in second-language grammar acquisition (Gallagher, Matsumoto, & Ohta, 2022), the causal role of the temporo-parietal cortex in novel word learning (Perceval, Martin, Copland, Laine, & Meinzer, 2017), and the domain-specificity of the dorsolateral prefrontal cortex in bilingual language control (Vaughn, Watlington, Linares Abrego, Tamber-Rosenau, & Hernandez, 2021). Transcranial alternating current stimulation (tACS) has also revealed important relationships, such as theta-phase synchronization in the frontoparietal regions causing visual memory matching (Polanía, Nitsche, Korman, Batsikadze, & Paulus, 2012), neural entrainment to speech causing intelligibility (Riecke, Formisano, Sorger, Başkent, & Gaudrain, 2018), and gamma activity in the temporal lobe causing moments of insight (Santarnecchi et al., 2019). Transcranial magnetic stimulation (TMS) has also been utilized to delineate functionally distinct language regions in the brain as well as to enhance performance in a variety of cognitive domains such as picture naming, numerical discrimination, and word recognition (Devlin & Watkins, 2007; Luber & Lisanby, 2014).

In dyslexia, three recent (systematic) reviews have already concluded that NIBS techniques are a promising remedial tool for reading deficits across age groups and languages, particularly emphasizing the efficacy of tDCS over the left temporo-parietal cortex (Cancer & Antonietti, 2018; Salehinejad, Ghanavati, Glinski, Hallajian, & Azarkolah, 2022; Turker & Hartwigsen, 2022). For example, several studies showed that stimulation (tDCS, TMS) over left temporo-parietal regions improved reading efficiency (speed and/or accuracy) (Costanzo, Menghini, Caltagirone, Oliveri, & Vicari, 2013; Costanzo et al., 2019, 2016; Turkeltaub et al., 2012). Although tACS studies were notably underrepresented, tACS was shown to improved phonemic awareness and phoneme categorization (Marchesotti et al., 2020; Rufener, Krauel, Meyer, Heinze, & Zaehle, 2019). Thus, various NIBS methods have already demonstrated their efficacy as a therapeutic intervention for various deficits in individuals with dyslexia. However, all of these stimulation methods suffer from a trade-off of focality and depth, which inherently limits their range of applicability and outright precludes the possibility of selectively stimulating the VWFA and deep-brain regions. It for that reason that we turn to tTIS.

### 4.2 Transcranial Temporal Interference Stimulation (tTIS)

Recently, a new method called tTIS has been developed (Grossman et al., 2017). To understand the utility of tTIS, it is helpful to understand the mechanisms of transcranial electrical stimulation methodologies used heretofore.

tDCS involves placing electrodes on the scalp and flowing a direct current from the anode(s) to the cathode(s) through a targeted region in the brain. In anodal stimulation, the current depolarizes neurons, increasing the probability of an action potential, thereby facilitating their activation during a cognitive task; in cathodal stimulation, neurons are hyperpolarized, reducing excitability and making action potentials harder to achieve (Nitsche et al., 2008). In tACS, an alternating current flows between electrodes, modulating the oscillatory activity of neuronal networks by entrainment, whereby neuronal firing rates align with the frequency of stimulation (Andrea Antal & Paulus, 2013). tACS can have either facilitatory or inhibitory effects depending on the frequency and relative phase of stimulation as well as stimulation site (Andrea Antal & Paulus, 2013). It is also important to note that the biochemical mechanisms involved in tES-induced plasticity remain to be fully elucidated (Fertonani & Miniussi, 2017).

Conventional tDCS and tACS rely on two large electrodes usually spaced far apart. Depending on the placement of electrodes, the stimulation can reach deep brain regions, however, this depth of penetration is accompanied by low focality, leading to stimulation of brain regions outside the region of interest, which additionally leads to greater interindividual variability (Mikkonen, Laakso, Tanaka, & Hirata, 2020). To solve the issue of focality, high-definition tES methods use a single small electrode surrounded by four small electrodes. This configuration allows for precise stimulation; however, it is limited to cortical areas (Thair, Holloway, Newport, & Smith, 2017). Thus, conventional tES and HD-tES methods pose a trade-off: You either have deep stimulation or focal stimulation, but not both. tTIS circumvents this trade-off by using two interfering high-frequency alternating currents.

tTIS relies on two key principles: First, high-frequency stimulation (> 1 kHz) has no effect on neural activity; second, overlapping electric fields of alternating currents interfere to create a beat frequency (Figure 1) at the difference of the two frequencies (Grossman et al., 2017). Therefore, the two electrode pairs and stimulation frequencies can be optimally chosen such that their electric fields overlap and interfere at any single given region in the brain without effectively stimulating surrounding regions. In this way, tTIS can achieve both focal and deep brain stimulation.

**Figure 1.**
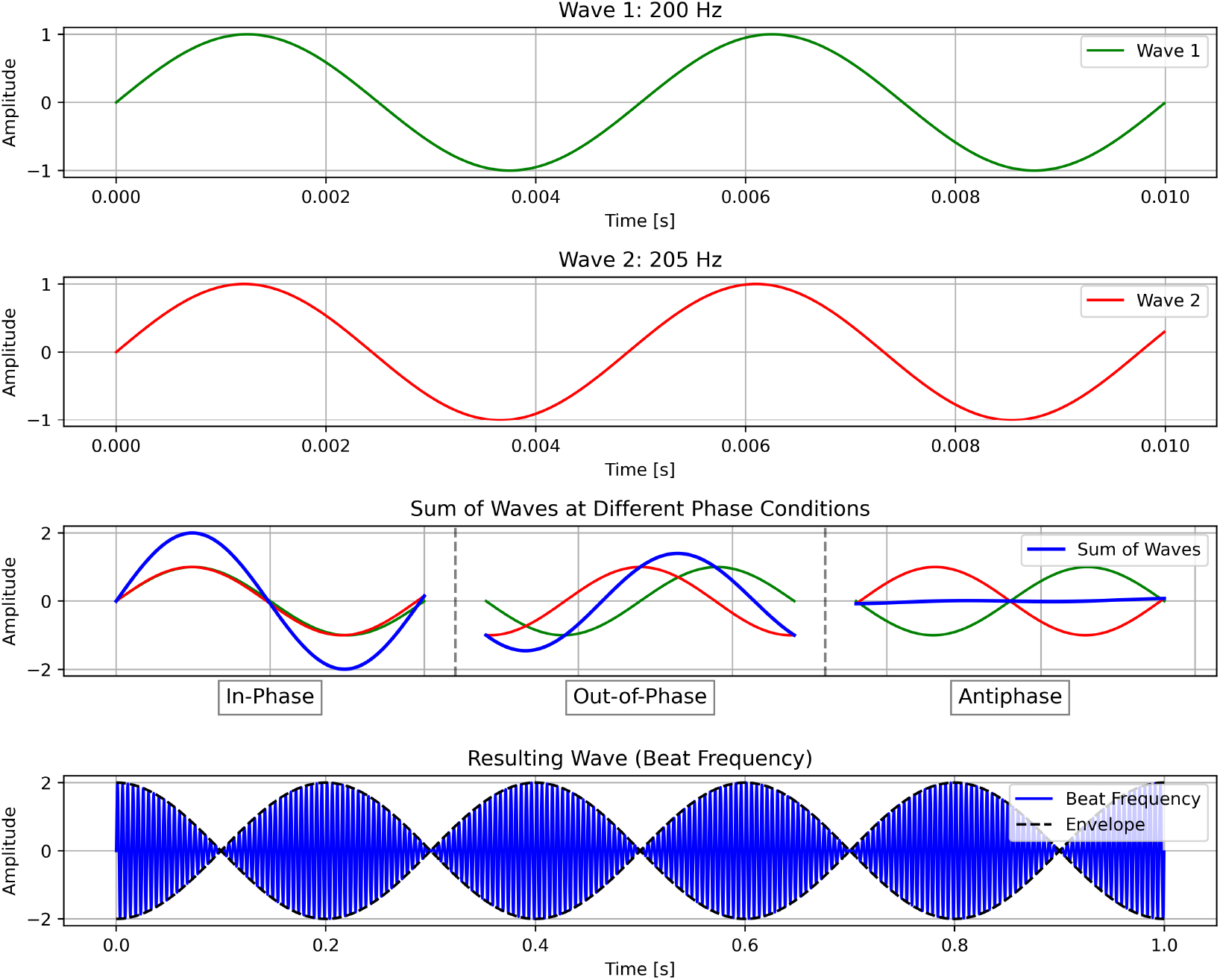
Schema of transcranial temporal interference stimulation (tTIS). In tTIS, high-frequency alternative currents interfere to create a beat frequency (Δf). For illustration purposes, this example shows 200 Hz and 205 Hz, although in practice, tTIS should utilize much higher frequencies (e.g., 2000 Hz).

As a recent development, tTIS has not yet been used in many studies investigating cognitive functions, including language. However, in one study, theta-burst tTIS of the striatum was shown to enhance motor learning in older healthy individuals (Wessel et al., 2023), which demonstrates the efficacy of tTIS to selectively stimulate deep-brain structures as well as the viability of using tTIS as a therapeutical intervention. To compare the efficacy of stimulation, we have simulated attempting to stimulate the VWFA with both HD-tDCS (Figure 2) and tTIS (Figure 3) using SimNIBS software (version 4.1.0) (Thielscher, Antunes, & Saturnino, 2015).

**Figure 2.**
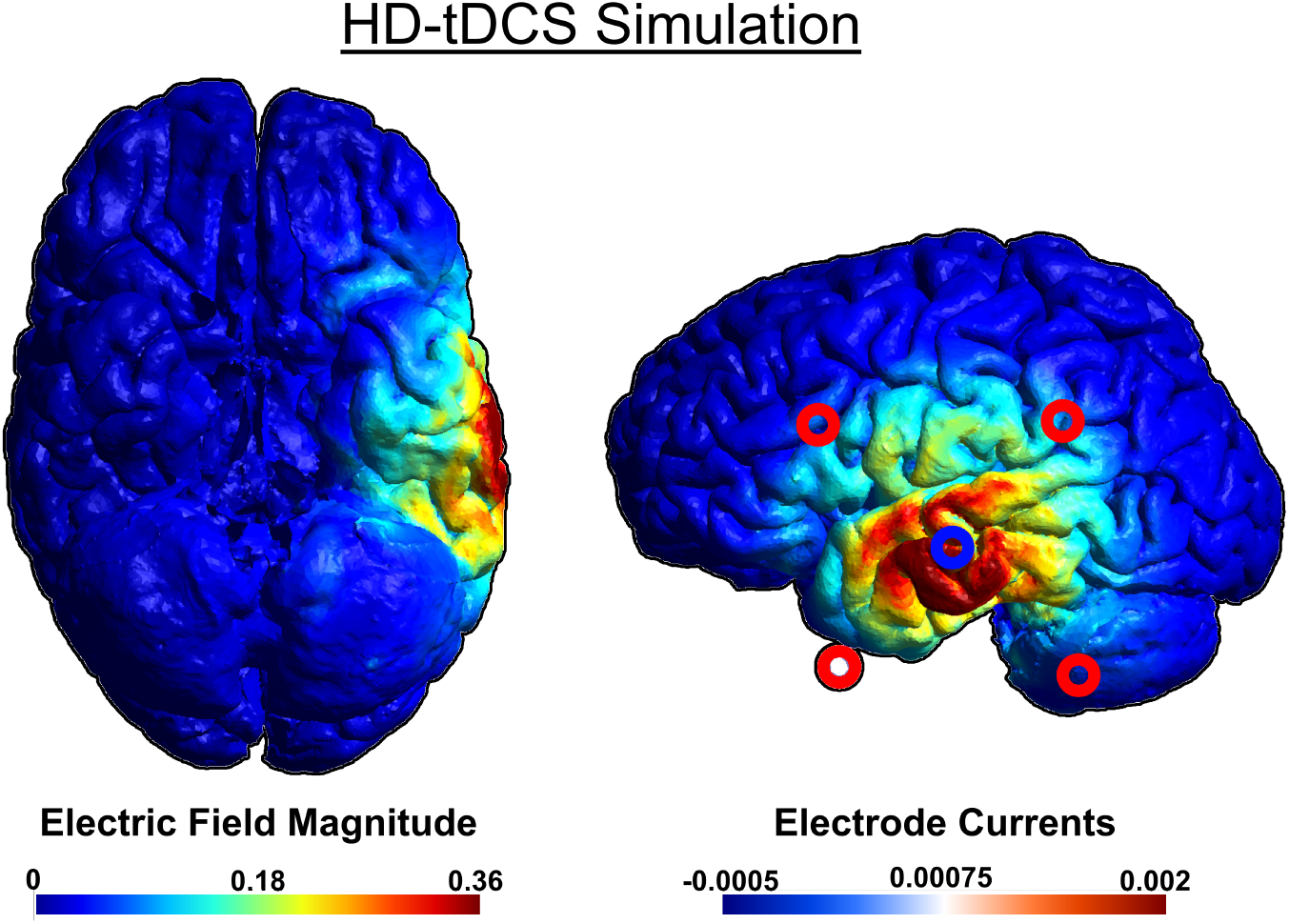
Simulation of HD-tDCS using a 4×1 ring electrode montage attempting to stimulate VWFA from the lowest available electrode positions. We used a central anode at PO10, with cathodes at O2, PO8, P8, and P10. For ease of visualization, stimulation electrodes were overlayed on the original image. Note: Because the scalp is not depicted in the rendering, the electrodes may appear to be floating due to perspective distortion when projecting a 3D image onto a 2D plane. This distortion affects the perceived distance between the electrodes and the brain surface.

**Figure 3.**
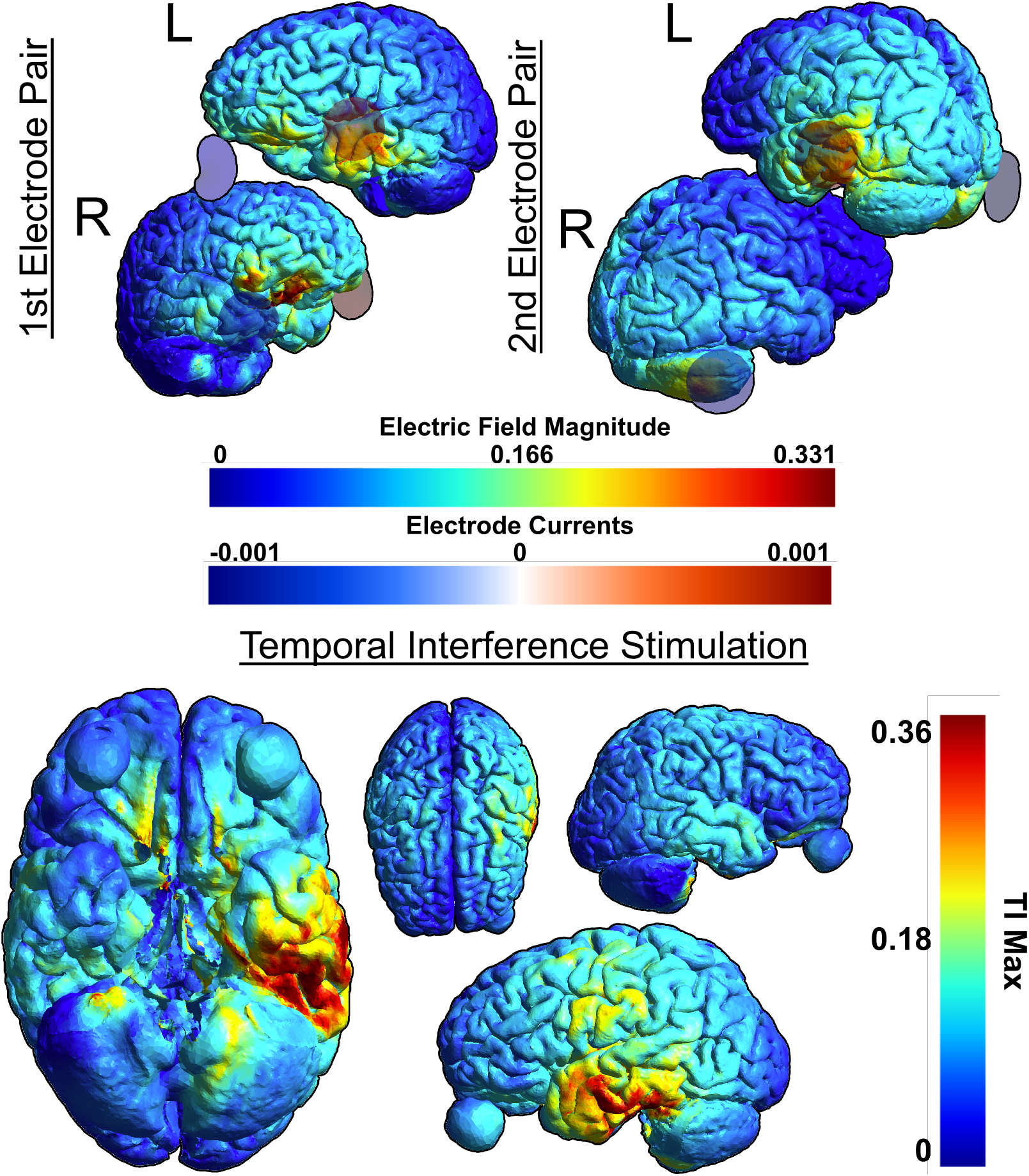
Simulation of tTIS on the VWFA. We used the TI Planning (TIP) tool by IT’IS software to compute the optimal electrode combination for stimulating the VWFA. The optimal electrode combinations were F7 and F10 and T7 and P8. We used the SimNIBS software for tTIS simulation. Note: Because the scalp is not depicted in the rendering, the electrodes may appear to be floating due to perspective distortion when projecting a 3D image onto a 2D plane. This distortion affects the perceived distance between the electrodes and the brain surface. Additionally, the 3D rendering in the lower panel includes the spheres representing the eyeballs.

It can be seen that while HD-tDCS fails to effectively stimulate the VWFA, which is difficult to reach on the ventral side of the occipito-temporal cortex, while tTIS can stimulate the VWFA much more effectively.

Although the mechanism of action of tTIS remains to be fully understood and it seems plausible to use tTIS for either downregulation or upregulation depending on stimulation parameters such as beat frequency, one study showed that tTIS downregulated neural activity (Carmona-Barrón et al., 2023), while previous stimulation studies failed to show facilitatory effects in already proficient readers (Cancer & Antonietti, 2018). Taking that together, we contend that it is more prudent to first attempt to use tTIS to elucidate the neurological pathogenesis of dyslexia subtypes and create a model for dyslexia subtypes in neurotypical adults. Thereafter, its use as novel therapeutical intervention for individuals with dyslexia can be investigated with a more comprehensive theoretical understanding.

Thus, by stimulating the brain regions revealed in a functional and structural clustering analysis and observing the behavioral effects of downregulation in those regions, we can infer relationships (whether they are ultimately causal or probabilistic) between neuropathologies and specific behavioral deficits of dyslexia. For example, if a clustering analysis reveals functional or structural anomalies in the VWFA, we can use tTIS to stimulate the VWFA and subsequently check for deficits in PA, RAN, etc. To ensure that any observed effects are specifically due to brain stimulation rather than placebo effects or unrelated variability, a sham stimulation group will serve as a control. Additionally, because dyslexia subtypes involve diverse cognitive deficits, it is crucial to test for a broad range of dyslexia-related symptoms rather than assuming a priori a one-to-one correspondence between the stimulated region and a single deficit. In this way, we can both reveal the neuropathologies of dyslexia subtypes and temporarily induce a specific subtype of dyslexia in neurotypical individuals for the purpose of investigating the efficacy of different therapeutic interventions.

Finally, it is important to note the safety of tTIS. Although it is a new method, it is mechanistically similar to tACS, thus tACS safety protocols and guidelines can be used as a baseline for considering the risks associated with tTIS. Although adverse effects such as headache, skin rash, fatigue, etc., are certainly possible, when following standard protocols, the likelihood of adverse effects is small, and their severity is generally mild (A. Antal et al., 2017; Matsumoto & Ugawa, 2017). Additionally, neurological effects induced by tACS are temporary (Kasten, Duecker, Maack, Meiser, & Herrmann, 2019), thus any induced dyslexic symptoms will dissipate within a few hours.

Taking all of this together, tTIS offers a safe method for downregulating activity at a single specific region anywhere in the brain, which, when applied judiciously, can temporarily create models of dyslexia subtypes in neurotypical adults.

## 5 Discussion

In this hypothesis article, we have proposed a dual-pronged approach for elucidating the neurological pathogenesis of distinctive dyslexia subtypes and hastening the development of therapeutical interventions for them. In the first prong, we propose a novel approach for analyzing functional and structural MRI data. Specifically, we suggest that by using PCA and clustering techniques, we can circumvent inadvertently grouping together different phenotypical expressions of dyslexia into a single study group. We can further reveal neuropathologically distinctive subgroups within the broader group of dyslexic individuals. This will inform the second prong of our approach, which revolves around the novel application of brain stimulation to dyslexia research. By using a novel tES method called tTIS, which allows for both deep brain and focal stimulation, we can selectively stimulate a specific region of the brain implicated in dyslexia and observe the behavioral deficits induced by the stimulation. This will allow the establishment of either causal or probabilistic relationships between the structurally/functionally anomalous brain regions involved in dyslexia and specific dyslexia symptoms, such as PA deficits, RAN deficits, etc. At the same time, by successfully inducing subtype-specific dyslexia symptoms, a model for each subtype can be created in neurotypical adults, which may in turn help aid our understanding of dyslexia from the perspective of single-, double-, or multiple-deficit perspectives.

As mentioned, the inability to use animal models that accurately reflect literacy deficits is a loss for dyslexia research. Using neurotypical adult human models for dyslexia would be a helpful substitute to fill in the gap. For example, if we have a reliable protocol for creating a dyslexia model whereby the neurotypical individual is induced with working memory-related perceptual deficits, we can more efficiently investigate and compare the efficacies of various behavioral interventions, such as working memory training protocols. Likewise, with a reliable model recreating PA deficits, we can efficiently assess behavioral interventions intended to help with PA deficits. In addition to behavioral interventions, pharmaceutical interventions may also be tested. For example, by giving a drug that is expected to upregulate activity in a given brain region, then subsequently using a stimulation protocol to downregulate activity in the same region, researchers can effectively compare the experimental group (i.e., drug plus stimulation) with the control group (i.e., stimulation only) to assess the efficacy of the drug. Using neurotypical models to investigate such pharmaceutical interventions could be more informative with greater statistical relevance than using an actual dyslexic group that consists of multiple disparate dyslexia subtypes. In this way, we can open the door to more efficiently investigate therapeutic interventions that are precisely tailored to the subtype of dyslexia afflicting the individual seeking treatment.

By no means do we hypothesize this to be a comprehensive solution, since it is inherently limited by its transient induction of symptoms and inability to morphologically replicate dyslexia subtypes, i.e., its inability to induce volumetric changes in stimulated brain regions. Indeed, a key limitation in our approach is that we are proposing to temporarily (i.e., short-term) replicate a developmental (i.e., long-term) disorder. Even aside from the aforementioned morphological differences, other features of the disorder may not translate perfectly to our model when compared to true dyslexia.

On the other hand, these limitations are clearly advantageous insofar as it would be both a grave violation of ethics and entirely undesirable to permanently impair a healthy participant’s cognitive function. Nevertheless, it stands to reason that certain structural abnormalities in dyslexia may not be properly investigated with this approach. Additionally, the approach outlined herein would not reveal anything of the genetic underpinnings of dyslexia or its subtypes. Thus, this approach is not intended to be one-size-fits-all kind of approach. However, from a functional perspective, we expect to adequately replicate the functional and connectivity deficits observed in dyslexia. To that end, certain aspects of dyslexia can be investigated more rapidly and more specifically than has been possible so far.

In sum, we propose that by combining a clustering approach to structural/functional MRI data with selective downregulating stimulation by tTIS, we can gain a deeper and more fundamental understanding of dyslexia subtypes, we can create subtype-specific dyslexia models in neurotypical adults, and we can ultimately improve the research environment for effective investigation into individually tailored treatment of dyslexia. Additionally, by demonstrating the efficacy of using tTIS to create neurotypical models of dyslexia, it may further prove the viability of employing the same methodological approach for investigating any number of other developmental and psychological disorders. Ultimately, we believe that by opening up this new avenue of research, we can more rapidly help improve the lives of those afflicted with dyslexia.

## 7 Author Contribution

DCG: Conceptualization, Writing - Original Draft, Writing - Review & Editing, Visualization, ZH: Conceptualization, Formal analysis, Writing - Original Draft, Writing - Review & Editing, Visualization, SO: Conceptualization, Writing - Review & Editing, Supervision, Project administration, and Funding acquisition.

## References

Antal, A., Alekseichuk, I., Bikson, M., Brockmöller, J., Brunoni, A. R., Chen, R., … Paulus, W. (2017). Low intensity transcranial electric stimulation: Safety, ethical, legal regulatory and application guidelines. Clinical Neurophysiology, 128(9), 1774–1809. 10.1016/j.clinph.2017.06.001

Antal, Andrea, & Paulus, W. (2013). Transcranial alternating current stimulation (tACS). Frontiers in Human Neuroscience, 7(JUN), 1–4. 10.3389/fnhum.2013.00317

Banfi, C., Koschutnig, K., Moll, K., Schulte-Körne, G., Fink, A., & Landerl, K. (2021). Reading-related functional activity in children with isolated spelling deficits and dyslexia. Language, Cognition and Neuroscience, 36(5), 543–561. 10.1080/23273798.2020.1859569

Beneventi, H., Tønnessen, F. E., & Ersland, L. (2009). Dyslexic Children Show Short-Term Memory Deficits in Phonological Storage and Serial Rehearsal: An fMRI Study. International Journal of Neuroscience, 119(11), 2017–2043. 10.1080/00207450903139671

Bergmann, T. O., & Hartwigsen, G. (2021). Inferring Causality from Noninvasive Brain Stimulation in Cognitive Neuroscience. Journal of Cognitive Neuroscience, 33(2), 195–225. 10.1162/jocn_a_01591

Boets, B., Op de Beeck, H. P., Vandermosten, M., Scott, S. K., Gillebert, C. R., Mantini, D., … Ghesquière, P. (2013). Intact But Less Accessible Phonetic Representations in Adults with Dyslexia. Science, 342(6163), 1251–1254. 10.1126/science.1244333

Bradley, L., & Bryant, P. E. (1978). Difficulties in auditory organisation as a possible cause of reading backwardness. Nature, 271(5647), 746–747. 10.1038/271746a0

Bruck, M. (1992). Persistence of Dyslexics’ Phonological Awareness Deficits. Developmental Psychology, 28(5), 874–886. 10.1037/0012-1649.28.5.874

Cancer, A., & Antonietti, A. (2018). tDCS Modulatory Effect on Reading Processes: A Review of Studies on Typical Readers and Individuals With Dyslexia. Frontiers in Behavioral Neuroscience, 12(July), 1–12. 10.3389/fnbeh.2018.00162

Carmona-Barrón, V. G., Fernández del Campo, I. S., Delgado-García, J. M., De la Fuente, A. J., Lopez, I. P., & Merchán, M. A. (2023). Comparing the effects of transcranial alternating current and temporal interference (tTIS) electric stimulation through whole-brain mapping of c-Fos immunoreactivity. Frontiers in Neuroanatomy, 17(March), 1–17. 10.3389/fnana.2023.1128193

Cattinelli, I., Borghese, N. A., Gallucci, M., & Paulesu, E. (2013). Reading the reading brain: A new meta-analysis of functional imaging data on reading. Journal of Neurolinguistics, 26(1), 214–238. 10.1016/j.jneuroling.2012.08.001

Cavalli, E., Chanoine, V., & Ziegler, J. C. (2023). MorphoSem. OpenNeuro, [Dataset]. 10.18112/openneuro.ds004786.v1.0.1

Centanni, T. M., Booker, A. B., Sloan, A. M., Chen, F., Maher, B. J., Carraway, R. S., … Kilgard, M. P. (2014). Knockdown of the dyslexia-associated gene Kiaa0319 impairs temporal responses to speech stimuli in rat primary auditory cortex. Cerebral Cortex, 24(7), 1753–1766. 10.1093/cercor/bht028

Centanni, Tracy Michelle, Booker, A. B., Chen, F., Sloan, A. M., Carraway, R. S., Rennaker, R. L., … Kilgard, M. P. (2016). Knockdown of dyslexia-gene Dcdc2 interferes with speech sound discrimination in continuous streams. Journal of Neuroscience, 36(17), 4895–4906. 10.1523/JNEUROSCI.4202-15.2016

Christodoulou, J. A., Del Tufo, S. N., Lymberis, J., Saxler, P. K., Ghosh, S. S., Triantafyllou, C., … Gabrieli, J. D. E. (2014). Brain Bases of Reading Fluency in Typical Reading and Impaired Fluency in Dyslexia. PLoS ONE, 9(7), e100552. 10.1371/journal.pone.0100552

Cohen, L., & Dehaene, S. (2004). Specialization within the ventral stream: the case for the visual word form area. NeuroImage, 22(1), 466–476. 10.1016/j.neuroimage.2003.12.049

Cohen, L., Dehaene, S., Naccache, L., Lehéricy, S., Dehaene-Lambertz, G., Hénaff, M.-A., & Michel, F. (2000). The visual word form area: Spatial and temporal characterization of an initial stage of reading in normal subjects and posterior split-brain patients. Brain, 123(2), 291–307. 10.1093/brain/123.2.291

Costanzo, F., Menghini, D., Caltagirone, C., Oliveri, M., & Vicari, S. (2013). How to improve reading skills in dyslexics: The effect of high frequency rTMS. Neuropsychologia, 51(14), 2953–2959. 10.1016/j.neuropsychologia.2013.10.018

Costanzo, F., Rossi, S., Varuzza, C., Varvara, P., Vicari, S., & Menghini, D. (2019). Long-lasting improvement following tDCS treatment combined with a training for reading in children and adolescents with dyslexia. Neuropsychologia, 130(January 2018), 38–43. 10.1016/j.neuropsychologia.2018.03.016

Costanzo, F., Varuzza, C., Rossi, S., Sdoia, S., Varvara, P., Oliveri, M., … Menghini, D. (2016). Reading changes in children and adolescents with dyslexia after transcranial direct current stimulation. NeuroReport, 27(5), 295–300. 10.1097/WNR.0000000000000536

Denckla, M. B., & Rudel, R. G. (1976). Rapid ‘automatized’ naming (R.A.N.): Dyslexia differentiated from other learning disabilities. Neuropsychologia, 14(4), 471–479. 10.1016/0028-3932(76)90075-0

Devlin, J. T., & Watkins, K. E. (2007). Stimulating language: Insights from TMS. Brain, 130(3), 610–622. 10.1093/brain/awl331

Fertonani, A., & Miniussi, C. (2017). Transcranial electrical stimulation: What we know and do not know about mechanisms. Neuroscientist, 23(2), 109–123. 10.1177/1073858416631966

Gallagher, D., Matsumoto, K., & Ohta, S. (2022). Causal evidence for the involvement of Broca’s area in second language acquisition : A longitudinal HD-tDCS study. BioRxiv. 10.1101/2022.12.19.520902 ;

Giraud, A.-L., & Poeppel, D. (2012). Cortical oscillations and speech processing: emerging computational principles and operations. Nature Neuroscience, 15(4), 511–517. 10.1038/nn.3063

Goto, M., Abe, O., Hagiwara, A., Fujita, S., Kamagata, K., Hori, M., … Daida, H. (2022). Advantages of Using Both Voxel- and Surface-based Morphometry in Cortical Morphology Analysis: A Review of Various Applications. Magnetic Resonance in Medical Sciences, 21(1), rev.2021-0096. 10.2463/mrms.rev.2021-0096

Grossman, N., Bono, D., Dedic, N., Kodandaramaiah, S. B., Rudenko, A., Suk, H.-J., … Boyden, E. S. (2017). Noninvasive Deep Brain Stimulation via Temporally Interfering Electric Fields. Cell, 169(6), 1029-1041.e16. 10.1016/j.cell.2017.05.024

Hasko, S., Bruder, J., Bartling, J., & Schulte-Körne, G. (2012). N300 indexes deficient integration of orthographic and phonological representations in children with dyslexia. Neuropsychologia, 50(5), 640–654. 10.1016/j.neuropsychologia.2012.01.001

He, Q., Xue, G., Chen, C., Chen, C., Lu, Z.-L., & Dong, Q. (2013). Decoding the Neuroanatomical Basis of Reading Ability: A Multivoxel Morphometric Study. Journal of Neuroscience, 33(31), 12835–12843. 10.1523/JNEUROSCI.0449-13.2013

Hynd, G. W., Hall, J., Novey, E. S., Eliopulos, D., Black, K., Gonzalez, J. J., … Cohen, M. (1995). Dyslexia and Corpus Callosum Morphology. Archives of Neurology, 52(1), 32–38. 10.1001/archneur.1995.00540250036010

Jednoróg, K., Gawron, N., Marchewka, A., Heim, S., & Grabowska, A. (2014). Cognitive subtypes of dyslexia are characterized by distinct patterns of grey matter volume. Brain Structure and Function, 219(5), 1697–1707. 10.1007/s00429-013-0595-6

Kasten, F. H., Duecker, K., Maack, M. C., Meiser, A., & Herrmann, C. S. (2019). Integrating electric field modeling and neuroimaging to explain inter-individual variability of tACS effects. Nature Communications, 10(1), 1–11. 10.1038/s41467-019-13417-6

Langer, N., Benjamin, C., Minas, J., & Gaab, N. (2015). The Neural Correlates of Reading Fluency Deficits in Children. Cerebral Cortex, 25(6), 1441–1453. 10.1093/cercor/bht330

Lazzaro, G., Fucà, E., Caciolo, C., Battisti, A., Costanzo, F., Varuzza, C., … Menghini, D. (2022). Understanding the Effects of Transcranial Electrical Stimulation in Numerical Cognition: A Systematic Review for Clinical Translation. Journal of Clinical Medicine, 11(8), 2082. 10.3390/jcm11082082

Lehongre, K., Morillon, B., Giraud, A. L., & Ramus, F. (2013). Impaired auditory sampling in dyslexia: Further evidence from combined fMRI and EEG. Frontiers in Human Neuroscience, 7(JUL), 1–8. 10.3389/fnhum.2013.00454

Lou, C., Duan, X., Altarelli, I., Sweeney, J. A., Ramus, F., & Zhao, J. (2019). White matter network connectivity deficits in developmental dyslexia. Human Brain Mapping, 40(2), 505–516. 10.1002/hbm.24390

Luber, B., & Lisanby, S. H. (2014). Enhancement of human cognitive performance using transcranial magnetic stimulation (TMS). NeuroImage, 85, 961–970. 10.1016/j.neuroimage.2013.06.007

Marchesotti, S., Nicolle, J., Merlet, I., Arnal, L. H., Donoghue, J. P., & Giraud, A.-L. (2020). Selective enhancement of low-gamma activity by tACS improves phonemic processing and reading accuracy in dyslexia. PLOS Biology, 18(9), e3000833. 10.1371/journal.pbio.3000833

Matsumoto, H., & Ugawa, Y. (2017). Adverse events of tDCS and tACS: A review. Clinical Neurophysiology Practice, 2, 19–25. 10.1016/j.cnp.2016.12.003

McGrath, L. M., Peterson, R. L., & Pennington, B. F. (2020). The Multiple Deficit Model: Progress, Problems, and Prospects. Scientific Studies of Reading, 24(1), 7–13. 10.1080/10888438.2019.1706180

McLaren, D. G., Ries, M. L., Xu, G., & Johnson, S. C. (2012). A generalized form of context-dependent psychophysiological interactions (gPPI): A comparison to standard approaches. NeuroImage, 61(4), 1277–1286. 10.1016/j.neuroimage.2012.03.068

Mikkonen, M., Laakso, I., Tanaka, S., & Hirata, A. (2020). Cost of focality in TDCS: Interindividual variability in electric fields. Brain Stimulation, 13(1), 117–124. 10.1016/j.brs.2019.09.017

Mukherjee, P., Roy, S., Ghosh, D., & Nandi, S. K. (2022). Role of animal models in biomedical research: a review. Laboratory Animal Research, 38(1), 1–17. 10.1186/s42826-022-00128-1

Nieto-Castanon, A., & Whitfield-Gabrieli, S. (2022). CONN functional connectivity toolbox: RRID SCR_009550, release 22. CONN functional connectivity toolbox: RRID SCR_009550, release 22. Hilbert Press. 10.56441/hilbertpress.2246.5840

Nitsche, M. A., Cohen, L. G., Wassermann, E. M., Priori, A., Lang, N., Antal, A., … Pascual-Leone, A. (2008). Transcranial direct current stimulation: State of the art 2008. Brain Stimulation, 1(3), 206–223. 10.1016/j.brs.2008.06.004

Norton, E. S., Beach, S. D., & Gabrieli, J. DE. (2015). Neurobiology of dyslexia. Current Opinion in Neurobiology, 30, 73–78. 10.1016/j.conb.2014.09.007

Norton, E. S., Black, J. M., Stanley, L. M., Tanaka, H., Gabrieli, J. D. E., Sawyer, C., & Hoeft, F. (2014). Functional neuroanatomical evidence for the double-deficit hypothesis of developmental dyslexia. Neuropsychologia, 61(650), 235–246. 10.1016/j.neuropsychologia.2014.06.015

Pennington, B. F. (2006). From single to multiple deficit models of developmental disorders. Cognition, 101(2), 385–413. 10.1016/j.cognition.2006.04.008

Pennington, B. F., Santerre-Lemmon, L., Rosenberg, J., MacDonald, B., Boada, R., Friend, A., … Olson, R. K. (2012). Individual prediction of dyslexia by single versus multiple deficit models. Journal of Abnormal Psychology, 121(1), 212–224. 10.1037/a0025823

Perceval, G., Martin, A. K., Copland, D. A., Laine, M., & Meinzer, M. (2017). High-definition tDCS of the temporo-parietal cortex enhances access to newly learned words. Scientific Reports, 7(1), 1–9. 10.1038/s41598-017-17279-0

Polanía, R., Nitsche, M. A., Korman, C., Batsikadze, G., & Paulus, W. (2012). The importance of timing in segregated theta phase-coupling for cognitive performance. Current Biology, 22(14), 1314–1318. 10.1016/j.cub.2012.05.021

Rack, J. P., Snowling, M. J., & Olson, R. K. (1992). The Nonword Reading Deficit in Developmental Dyslexia: A Review. Reading Research Quarterly, 27(1), 28. 10.2307/747832

Richlan, F., Kronbichler, M., & Wimmer, H. (2009). Functional abnormalities in the dyslexic brain: A quantitative meta-analysis of neuroimaging studies. Human Brain Mapping, 30(10), 3299–3308. 10.1002/hbm.20752

Richlan, F., Kronbichler, M., & Wimmer, H. (2011). Meta-analyzing brain dysfunctions in dyslexic children and adults. NeuroImage, 56(3), 1735–1742. 10.1016/j.neuroimage.2011.02.040

Richlan, F., Kronbichler, M., & Wimmer, H. (2013). Structural abnormalities in the dyslexic brain: A meta-analysis of voxel-based morphometry studies. Human Brain Mapping, 34(11), 3055–3065. 10.1002/hbm.22127

Riecke, L., Formisano, E., Sorger, B., Başkent, D., & Gaudrain, E. (2018). Neural Entrainment to Speech Modulates Speech Intelligibility. Current Biology, 28(2), 161-169.e5. 10.1016/j.cub.2017.11.033

Robinson, N. B., Krieger, K., Khan, F., Huffman, W., Chang, M., Naik, A., … Gaudino, M. (2019). The current state of animal models in research: A review. International Journal of Surgery, 72(August), 9–13. 10.1016/j.ijsu.2019.10.015

Roitsch, J., & Watson, S. (2019). An Overview of Dyslexia: Definition, Characteristics, Assessment, Identification, and Intervention. Science Journal of Education, 7(4), 81. 10.11648/j.sjedu.20190704.11

Rufener, K. S., Krauel, K., Meyer, M., Heinze, H. J., & Zaehle, T. (2019). Transcranial electrical stimulation improves phoneme processing in developmental dyslexia. Brain Stimulation, 12(4), 930–937. 10.1016/j.brs.2019.02.007

Russell, V. A. (2011). Overview of Animal Models of Attention Deficit Hyperactivity Disorder (ADHD). Current Protocols in Neuroscience, 54(1), 1–25. 10.1002/0471142301.ns0935s54

Sagvolden, T., Metzger, M. A., Schiorbeck, H. K., Rugland, A.-L., Spinnangr, I., & Sagvolden, G. (1992). The spontaneously hypertensive rat (SHR) as an animal model of childhood hyperactivity (ADHD): changed reactivity to reinforcers and to psychomotor stimulants. Behavioral and Neural Biology, 58(2), 103–112. 10.1016/0163-1047(92)90315-U

Salehinejad, M. A., Ghanavati, E., Glinski, B., Hallajian, A., & Azarkolah, A. (2022). A systematic review of randomized controlled trials on efficacy and safety of transcranial direct current stimulation in major neurodevelopmental disorders: ADHD, autism, and dyslexia. Brain and Behavior, 12(9), 1–21. 10.1002/brb3.2724

Santarnecchi, E., Sprugnoli, G., Bricolo, E., Costantini, G., Liew, S.-L., Musaeus, C. S., … Rossi, S. (2019). Gamma tACS over the temporal lobe increases the occurrence of Eureka! moments. Scientific Reports, 9(1), 5778. 10.1038/s41598-019-42192-z

Snowling, M. (1998). Dyslexia as a Phonological Deficit: Evidence and Implications. Child Psychology and Psychiatry Review, 3(1), 4–11. 10.1017/S1360641797001366

Snowling, M. J. (1981). Phonemic deficits in developmental dyslexia. Psychological Research, 43(2), 219–234. 10.1007/BF00309831

Sokhadze, E. M., El-Baz, A. S., Tasman, A., Sears, L. L., Wang, Y., Lamina, E. V., & Casanova, M. F. (2014). Neuromodulation Integrating rTMS and Neurofeedback for the Treatment of Autism Spectrum Disorder: An Exploratory Study. Applied Psychophysiology and Biofeedback, 39(3–4), 237–257. 10.1007/s10484-014-9264-7

Stoyanov, D., Kandilarova, S., Paunova, R., Barranco Garcia, J., Latypova, A., & Kherif, F. (2019). Cross-Validation of Functional MRI and Paranoid-Depressive Scale: Results From Multivariate Analysis. Frontiers in Psychiatry, 10(November), 1–8. 10.3389/fpsyt.2019.00869

Thair, H., Holloway, A. L., Newport, R., & Smith, A. D. (2017). Transcranial Direct Current Stimulation (tDCS): A Beginner’s Guide for Design and Implementation. Frontiers in Neuroscience, 11(NOV). 10.3389/fnins.2017.00641

Thielscher, A., Antunes, A., & Saturnino, G. B. (2015). Field modeling for transcranial magnetic stimulation: A useful tool to understand the physiological effects of TMS? Proceedings of the Annual International Conference of the IEEE Engineering in Medicine and Biology Society, EMBS, 2015-Novem, 222–225. 10.1109/EMBC.2015.7318340

Turkeltaub, P. E., Benson, J., Hamilton, R. H., Datta, A., Bikson, M., & Coslett, H. B. (2012). Left lateralizing transcranial direct current stimulation improves reading efficiency. Brain Stimulation, 5(3), 201–207. 10.1016/j.brs.2011.04.002

Turker, S., & Hartwigsen, G. (2022). The use of noninvasive brain stimulation techniques to improve reading difficulties in dyslexia: A systematic review. Human Brain Mapping, 43(3), 1157–1173. 10.1002/hbm.25700

Turker, S., Kuhnke, P., Jiang, Z., & Hartwigsen, G. (2023). Disrupted network interactions serve as a neural marker of dyslexia. Communications Biology, 6(1), 1114. 10.1038/s42003-023-05499-2

Van Boxtel, G. J. M. (1998). Computational and statistical methods for analyzing event-related potential data. Behavior Research Methods, Instruments, & Computers, 30(1), 87–102. 10.3758/BF03209419

Varghese, M., Keshav, N., Jacot-Descombes, S., Warda, T., Wicinski, B., Dickstein, D. L., … Hof, P. R. (2017). Autism spectrum disorder: neuropathology and animal models. Acta Neuropathologica, 134(4), 537–566. 10.1007/s00401-017-1736-4

Vaughn, K. A., Watlington, E. M., Linares Abrego, P., Tamber-Rosenau, B. J., & Hernandez, A. E. (2021). Prefrontal transcranial direct current stimulation (tDCS) has a domain-specific impact on bilingual language control. Journal of Experimental Psychology: General, 150(5), 996–1007. 10.1037/xge0000956

Wessel, M. J., Beanato, E., Popa, T., Windel, F., Vassiliadis, P., Menoud, P., … Hummel, F. C. (2023). Noninvasive theta-burst stimulation of the human striatum enhances striatal activity and motor skill learning. Nature Neuroscience, 26(11), 2005–2016. 10.1038/s41593-023-01457-7

Williams, V. J., Juranek, J., Cirino, P., & Fletcher, J. M. (2018). Cortical Thickness and Local Gyrification in Children with Developmental Dyslexia. Cerebral Cortex, 28(3), 963–973. 10.1093/cercor/bhx001

Wolf, M., & Bowers, P. G. (1999). The double-deficit hypothesis for the developmental dyslexias. Journal of Educational Psychology, 91(3), 415–438. 10.1037/0022-0663.91.3.415

Yang, L.-P., Li, C.-B., Li, X.-M., Zhai, M.-M., Zhao, J., & Weng, X.-C. (2022). Prevalence of developmental dyslexia in primary school children: a protocol for systematic review and meta-analysis. World Journal of Pediatrics, 18(12), 804–809. 10.1007/s12519-022-00572-y

Žaric, G., Timmers, I., Gerretsen, P., Fraga González, G., Tijms, J., van der Molen, M. W., … Bonte, M. (2018). Atypical White Matter Connectivity in Dyslexic Readers of a Fairly Transparent Orthography. Frontiers in Psychology, 9(JUN), 1–15. 10.3389/fpsyg.2018.01147

