## Supplementary Materials for "Modeling Dyslexia in Neurotypical Adults by Combining Neuroimaging and Neuromodulation Techniques: A Hypothesis Paper"

### Preliminary Structural Analysis

In a preliminary analysis, we used participants from two MRI studies. The first dataset included 22 typically developing children (age: 9.05 ± 0.69 years) and 20 children with dyslexia (age: 9.05 ± 0.69 years), consisting of 11 males and 11 females in the typically developing group, and 13 males and 7 females in the dyslexia group. All participants were German speakers (Banfi et al., 2021). The second dataset comprised 22 typical adult readers (age: 22.7 ± 4.2 years) and 20 adults with dyslexia (age: 22.7 ± 4.2 years), with 12 males and 10 females in the typical readers group, and 13 males and 7 females in the dyslexia group. These participants were French speakers (Cavalli et al., 2023). Both datasets contained structural and functional MRI data, however, thus far, only structural data have been analyzed.

While the datasets were selected based on open-source availability, we recognize that differences in language orthographic depth (German vs. French) and age (children vs. adults) may limit the comparability of results. The selection of these datasets was primarily driven by the need to explore structural and functional variations in dyslexia, with a focus on leveraging existing, well-characterized MRI data.

Statistical Parametric Mapping (SPM12) software (Friston et al., 1994) on MATLAB R2024a was used along with the CAT12 toolbox (Gaser et al., 2024) to perform Voxel-based Morphometry (VBM) / Surface-based Morphometry (SBM) and region of interest (ROI) analysis. VBM analysis focuses on the 3D volume of brain tissue and is primarily used for analyzing gray and white matter volumes across brain regions. The preprocessing steps of the structural MRI data included brain tissue segmentation into the cerebrospinal fluid (CSF), white matter (WM), and gray matter (GM) using the SPM default template, and data smoothing for signal averaging and noise cancelation. At the same time, SBM works with the cortical surface, analyzing features such as cortical thickness, sulcal depth, and gyrification, and is often used to study more localized cortical regions. The preprocessing steps included cortical thickness and central surface estimation, topology correction, spherical mapping, and spherical registration. We used the "Estimate Mean Values inside ROI" function in CAT12 GUI to extract the ROI data, such as gray matter volumes. We used the DK40 atlas (Desikan et al., 2006) and defined the right and left FG as our ROIs.

Through voxel-based morphometry, we found significant structural differences between dyslexic individuals and controls (corrected $p < 0.05$) (Supplementary Table 1). Dyslexic adults exhibited widespread reductions in white matter volume, particularly in the left corpus callosum, which may impair interhemispheric communication and influence the right hemisphere's functions. Significant reductions were observed in key tracts such as the left arcuate fasciculus and right inferior fronto-occipital fasciculus, which are crucial for language processing and visual-spatial integration, respectively.

**Supplementary Table 1:** Locations of white matter volume reductions in dyslexic patients.

| Brain regions | Corrected P | T | MNI coordinates | Cluster size (mm$^3$) |
|---|---|---|---|---|
| L. Corpus Callosum | 0.001 | 4.70 | –15, –28, 38 | 1791 |
| R. inferior fronto-occipital fasciculus | 0.001 | 4.60 | 34, 28, 9 | 1560 |
| L. Corpus Callosum | 0.008 | 4.44 | –15, 24, 33 | 979 |
| R. Cingulum Bundle | 0.038 | 4.67 | 12, -57, 40 | 598 |
| L. Arcuate Fasciculus (post. Segment) | 0.035 | 4.89 | –48, –36, 26 | 652 |

We used cluster-based false discovery rate (FDR) correction.

Moreover, dyslexic adults showed cortical thinning in the right FG and rostral anterior cingulate cortex (rACC), regions linked to visual word recognition and cognitive control, respectively (corrected $p < 0.05$) (Supplementary Figure 1).

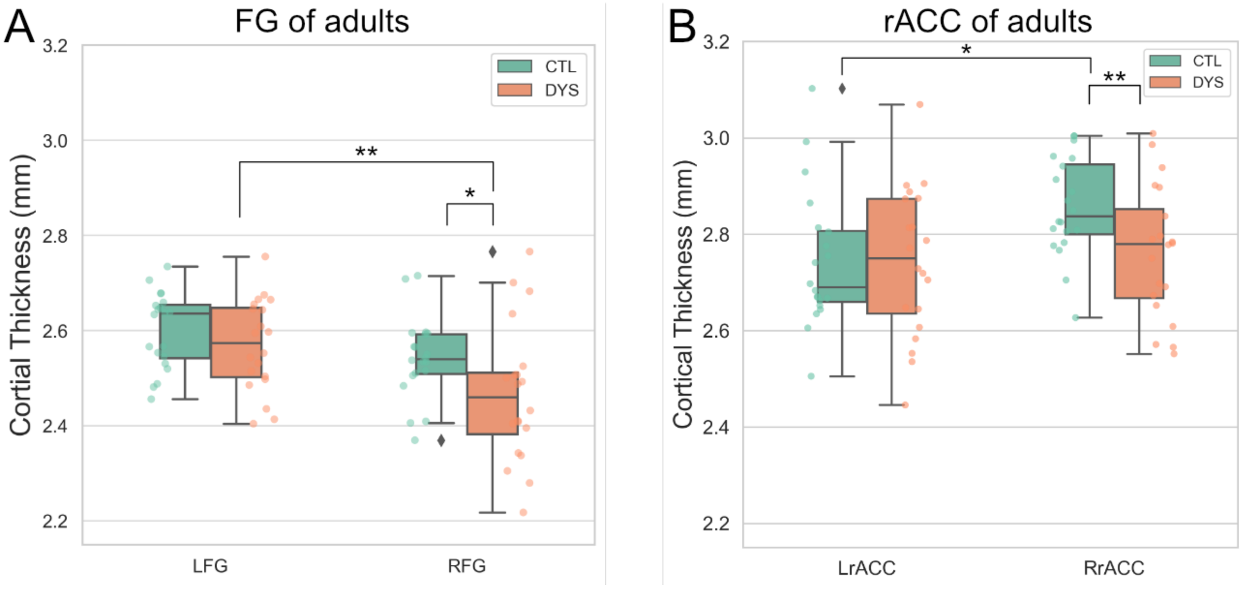


**Supplementary Figure 1.** Cortical thickness differences across groups. Results of two-way analysis of variance (ANOVA) (Group [Controls vs. Dyslexics] × Hemisphere [Left vs. Right]). **A**) left and right fusiform gyrus (FG) and **B**) rostral anterior cingulate cortex (rACC). * corrected $p < .05$, ** corrected $p < .01$. CTL: Controls, DYS: Dyslexics.

Dyslexic individuals further showed a distinct aging-related pattern of cortical thinning in the right FG and rACC, with greater thinning observed in older adults with dyslexia compared to controls (corrected $p < 0.05$) (Supplementary Figure 2).

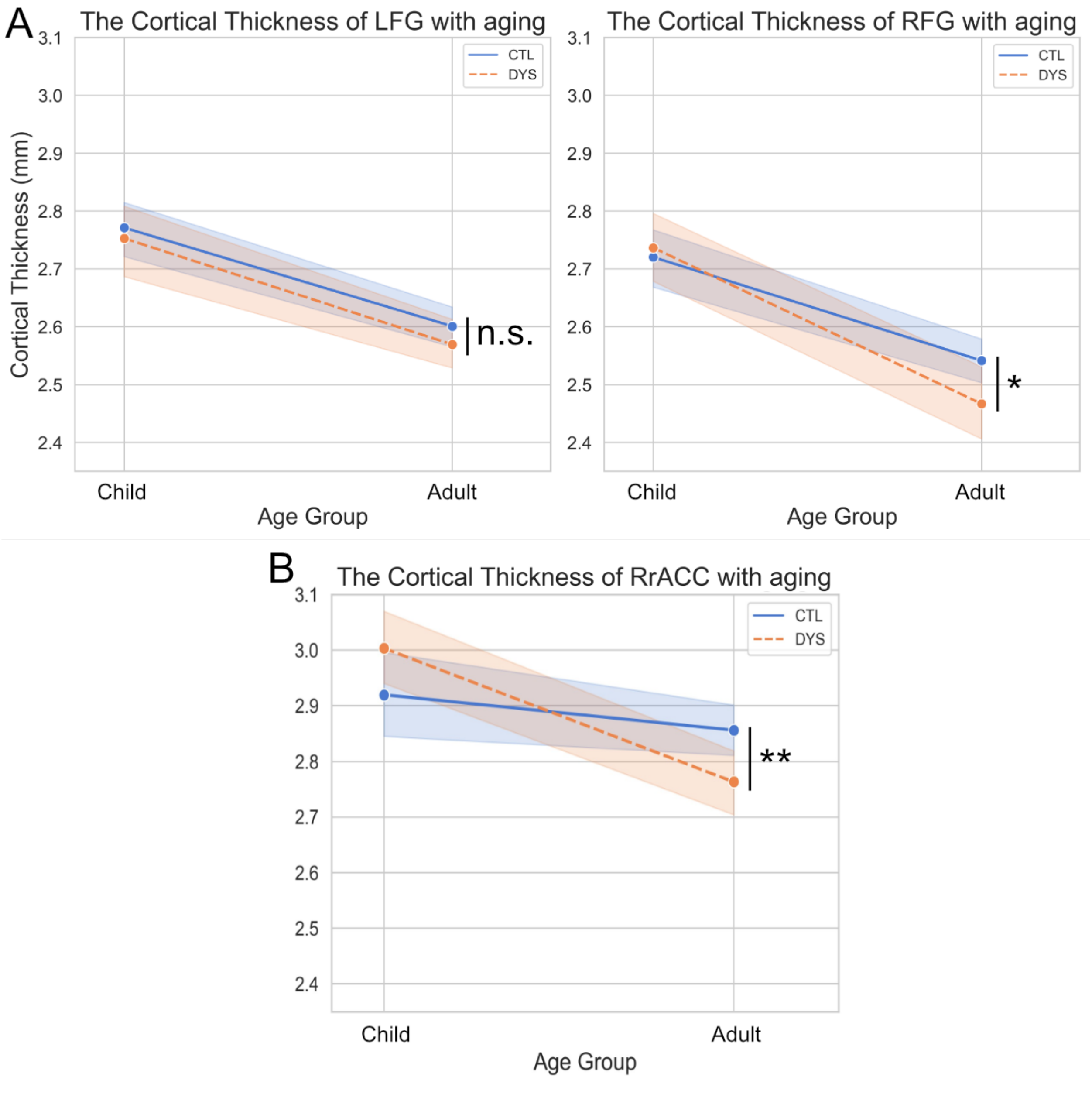


**Supplementary Figure 2.** Cortical thickness changes by aging. Results of two-way ANOVA (Group [Controls vs. Dyslexics] × Age [Children vs. Adults]). **A**) left and right FG and **B**) right rACC. * corrected $p < .05$, ** corrected $p < .01$, n.s. = not significant. CTL: Controls, DYS: Dyslexics.

These findings suggest that the structural deficits observed in dyslexia are not limited to the left hemisphere but extend to the right hemisphere, likely because of compensatory mechanisms. The reduction in the corpus callosum may disrupt inter-hemispheric communication, contributing to the observed thinning in the right hemisphere. Furthermore, the thinner cortex in the right FG and rACC may indicate impaired function linked to reduced connectivity. These structural abnormalities are likely to underlie the reading and cognitive challenges faced by individuals with dyslexia.

However, because brain stimulation modulates activity, structural results alone do not provide a robust or reliable basis for defining a stimulation target. Therefore, combining structural and functional MRI results to identify areas of alignment between the two will provide more valuable insights, which we will address in future analyses.
